# Cloning of Nine Glucocorticoid Receptor Isoforms from the Slender African lungfish (*Protopterus dolloi*)

**DOI:** 10.1101/2022.03.19.484994

**Authors:** Yoshinao Katsu, Shin Oana, Xiaozhi Lin, Susumu Hyodo, Laurent Bianchetti, Michael E. Baker

## Abstract

We wanted to clone the glucocorticoid receptor (GR) from slender African lungfish (*Protopterus dolloi*) for comparison to *P. dolloi* MR, which we had cloned and were characterizing, as well as for comparison to the GRs from humans, elephant shark and zebrafish. However, although sequencing of the genome of the Australian lungfish *(Neoceratodus forsteri)*, as well as, that of the West African lungfish (*Protopterus annectens*) were reported in the first three months of 2021, we could not retrieve a GR sequence with a BLAST search of GenBank, when we submitted our research for publication in July 2021. Moreover, we were unsuccessful in cloning the GR from slender African lungfish using a cDNA from the ovary of *P. dolloi* and PCR primers that had successfully cloned a GR from elephant shark, *Xenopus* and gar GRs. On October 21, 2021 the nucleotide sequence of West African lungfish (*P. annectens*) GR was deposited in GenBank. We used this GR sequence to construct PCR primers that successfully cloned the GR from the slender spotted lungfish. Here, we report the sequences of nine *P. dolloi* GR isoforms and explain the basis for the previous failure to clone a GR from slender African lungfish using PCR primers that cloned the GR from elephant shark, *Xenopus* and gar. Studies are underway to determine corticosteroid activation of these slender African lungfish GRs.

## 1. INTRODUCTION

The glucocorticoid receptor (GR) belongs to the nuclear receptor family, a diverse group of transcription factors that arose in multicellular animals [1–4]. The GR has many key roles in the physiology of humans and other terrestrial vertebrates and fish [5–8]. Important for understanding the function of the GR is that it is closely related to the mineralocorticoid receptor (MR) [9–11]. These two steroid receptors evolved from a duplication of an ancestral corticoid receptor (CR) in a jawless fish (cyclostome), which has descendants in modern lampreys and hagfish [11–13]. A distinct GR and MR first appear in cartilaginous fishes (Chondrichthyes) [1,9,11,14,15], which diverged from bony vertebrates about 450 million years ago [16,17].

Lungfish are important in the transition of vertebrates from water to land [18–22], and aldosterone activation of the MR is important in this process [11,22–25]. Aldosterone, the main physiological mineralocorticoid in humans and other terrestrial vertebrates [26–29], first appears in lungfish [21–23]. To investigate the origins of aldosterone signaling, we cloned the MR from slender spotted African lungfish (*P. dolloi*) and studied its activation by aldosterone, other corticosteroids and progesterone [30]. To continue our investigation of early events in the evolution of the GR and MR, we sought to clone the *P. dolloi* GR for comparison with *P. dolloi* MR, as well as with the GR in coelacanths, zebrafish and humans. However, a BLAST search with the sequence of the GR from coelacanth and zebrafish did not retrieve the sequence of *P. dolloi* GR or any other lungfish GR from GenBank. Nor could we clone the *P. dolloi* GR using a cDNA from *P. dolloi* ovary using PCR primers that had successfully cloned a GR from elephant shark GR [15] and chicken, alligator and frog GRs [31]. Fortunately, on October 21, 2021 the nucleotide sequence of African lungfish (*P. annectens*) GR was deposited in GenBank, which gave us sufficient information for PCR primers to clone nine isoforms of *P. dolloi* GR. Here we report the sequences of these nine *P. dolloi* GR isoforms and explain the basis for the previous failure to clone a GR from slender African lungfish using PCR primers that previously cloned the GR from elephant shark, *Xenopus* and gar [15,31,32]. Our analysis of these nine GR sequences indicates that they evolved by alternative splicing and gene duplication [33,34].

## 2. Results and Discussion

### 2A. Multiple sequence alignment of nine *P. dolloi* GR isoforms

Figure 1 shows a multiple sequence alignment of the nine isoforms of *P. dolloi* GR. The nine *P. dolloi* GRs cluster into three groups: group I (GR-A1, GR-A2), group II (GR-B1, GR-B2, GR-B3) and group III (GR-C1, GR-C2, GR-C3, GR-C4). GR-A2 begins at “MMDP”, a sequence motif that is conserved in all nine GRs.

**Figure 1A and 1B.**
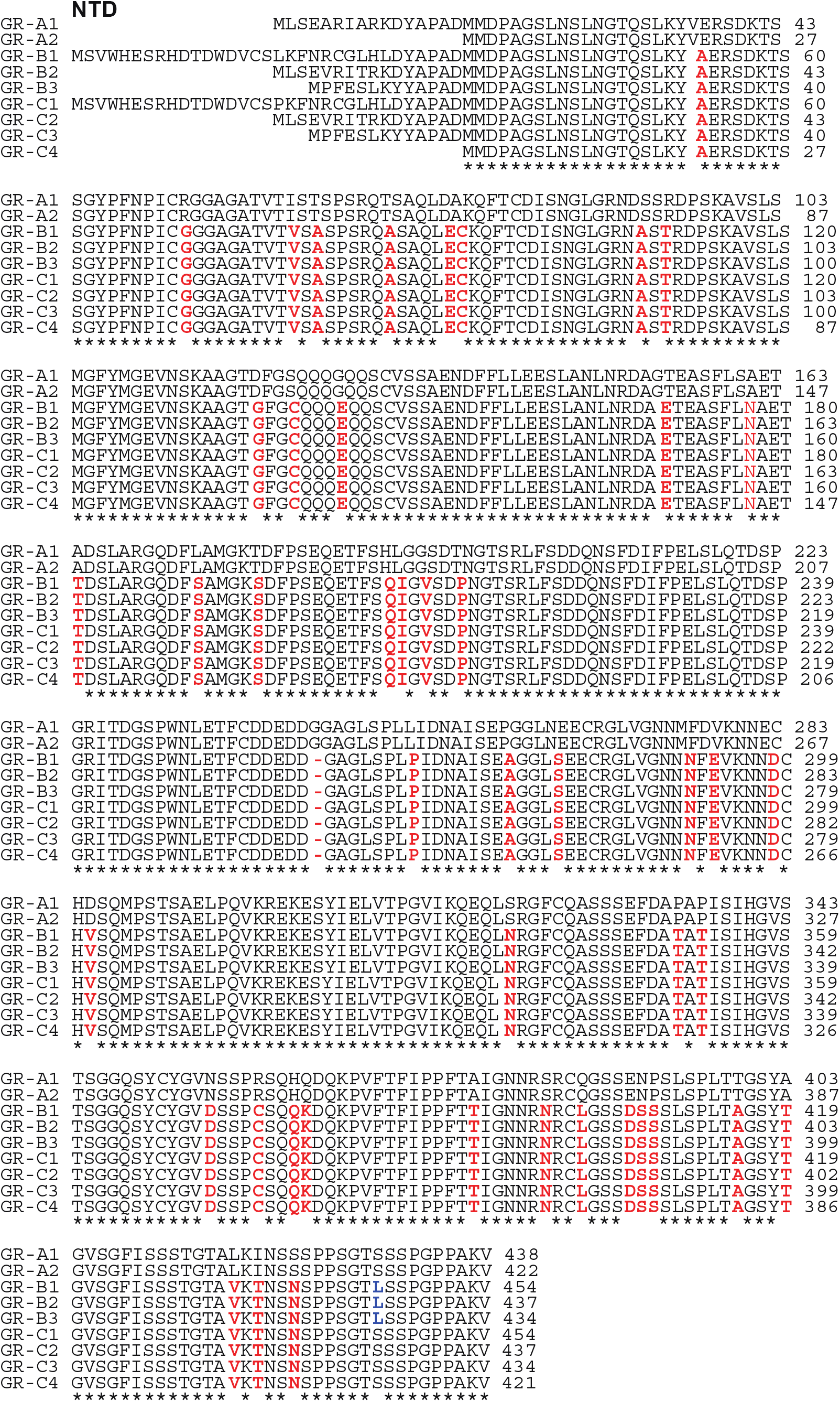
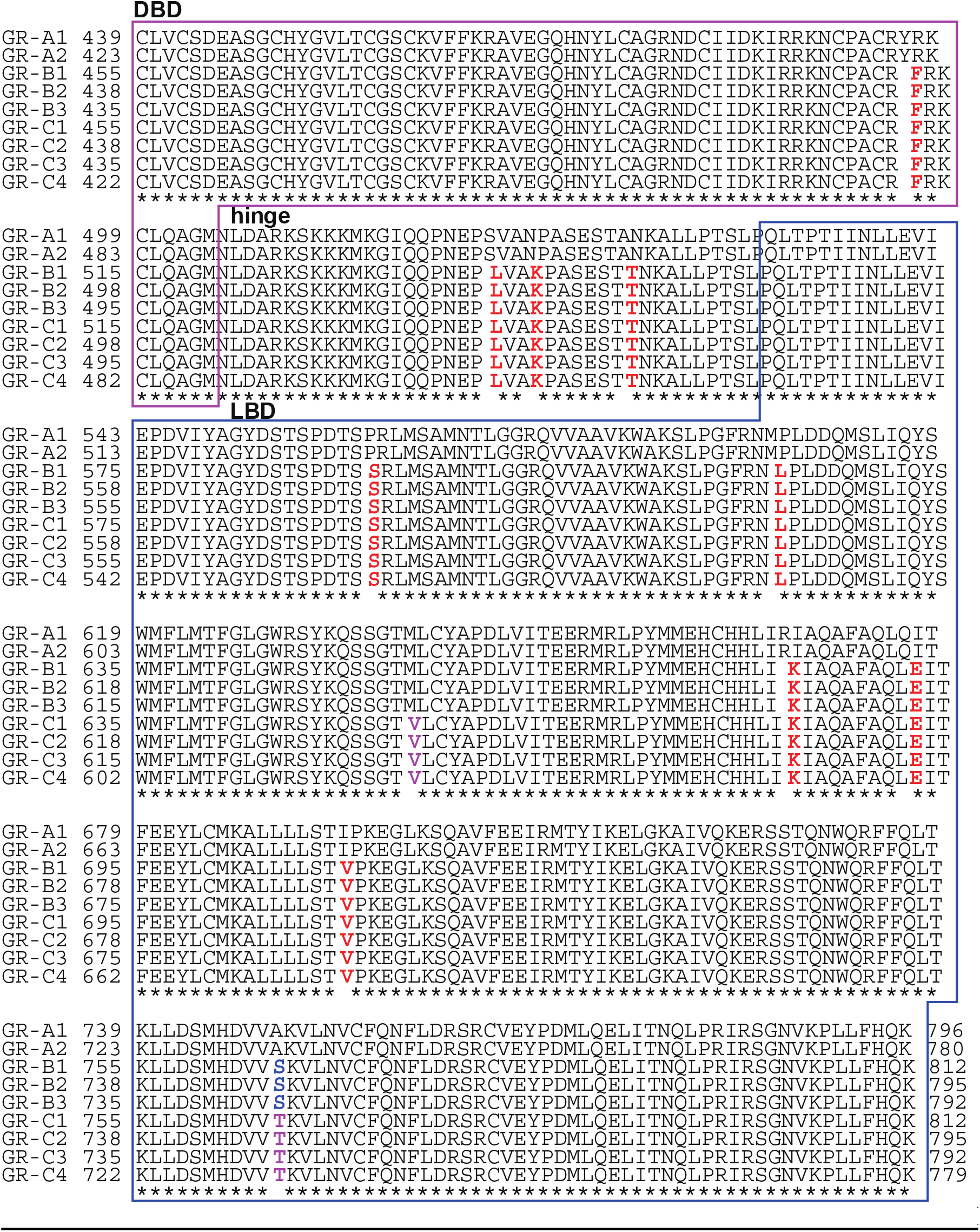
Multiple Alignment of the amino acid sequences slender African lungfish glucocorticoid receptors. Total RNA was isolated from *P. dolloi* ovary and translated into cDNA. PCR was performed using four primer sets based on the sequence of *P. annectens* GR, as described in the Methods section. The amplified DNA fragments were sub-cloned into a vector for sequence analysis. Similar to other steroid receptors, slender African lungfish GR can be divided into four functional domains [6,8], consisting of a ligand-binding domain (LBD) at the C-terminus, a DNA-binding domain (DBD) in the center that is joined to the LBD by a short hinge domain (hinge), and a domain at the amino-terminus (NTD). GenBank accession no. BDF84376 for GR-A1, BDF84377 for GR-A2, BDF84378 for GR-B1, BDF84379 for GR-B2, BDF84380 for GR-B3, BDF84381 for GR-C1, BDF84382 for GR-C2, BDF84383 for GR-C3, and BDF84384 for GR-C4. Sequences were aligned with Clustal W [35], as described in the Methods section.

The multiple alignment reveals that these nine slender African lungfish GRs evolved through alternative splicing and gene duplications (Figure 1). GR-A2 appears to be a product of alternative splicing of GR-A1. GR-C4 appears to be a product of alternative splicing of one or more GR-C isoforms, which supports a GR gene duplication in *P. dolloi* genome. There also is evidence for gene duplications among the *P. dolloi* GRs. MLSE at the beginning of GR-A1 is conserved in GR-B2 and GR-C2. A closely following YAPAD sequence is conserved in all *P. dolloi* GR isoforms. Fifteen of the first sixteen amino acids at the amino terminus of GR-A-1 are conserved in GR-B2 and GR-C2 (Figure 1A). This amino acid sequence is highly conserved in the other seven GRs. The rest of GR-A2 beginning at MMDPAGALNSLNGTQSLNKY is identical in GR-A1, and this amino acid sequence is highly conserved in the other seven GRs. MPFESLKYYAPAD is conserved at the beginning of GR-B3 and GR-C3. Beginning at the conserved MMDP sequence in the N-terminal domain, the two GR-A isoforms differ at 55 positions from the three GR-B and the four GR-C isoforms.

### 2B. Comparison of slender African lungfish GRs and West African lungfish GRs

To begin to understand sequence conservation and divergence among lungfish GRs, we compared GR-A1, GR-B1 and GR-C1, which are the three longest slender African lungfish GRs, with the four West African lungfish glucocorticoid receptor sequences in GenBank (Figure 2). The multiple sequence alignment, shown in Figure 2, reveals strong sequence conservation in the DBD, with a difference at only one position containing a semi-conserved phenylalanine-tyrosine. The sequences in the LBD and hinge domains of slender African lungfish GR and West African lungfish GR also are highly conserved. There are small segments of sequence divergence in the NTD, but most of the NTD is conserved. Overall slender African lungfish GRs and African lungfish GRs are very similar to each other.

**Figure 2.**
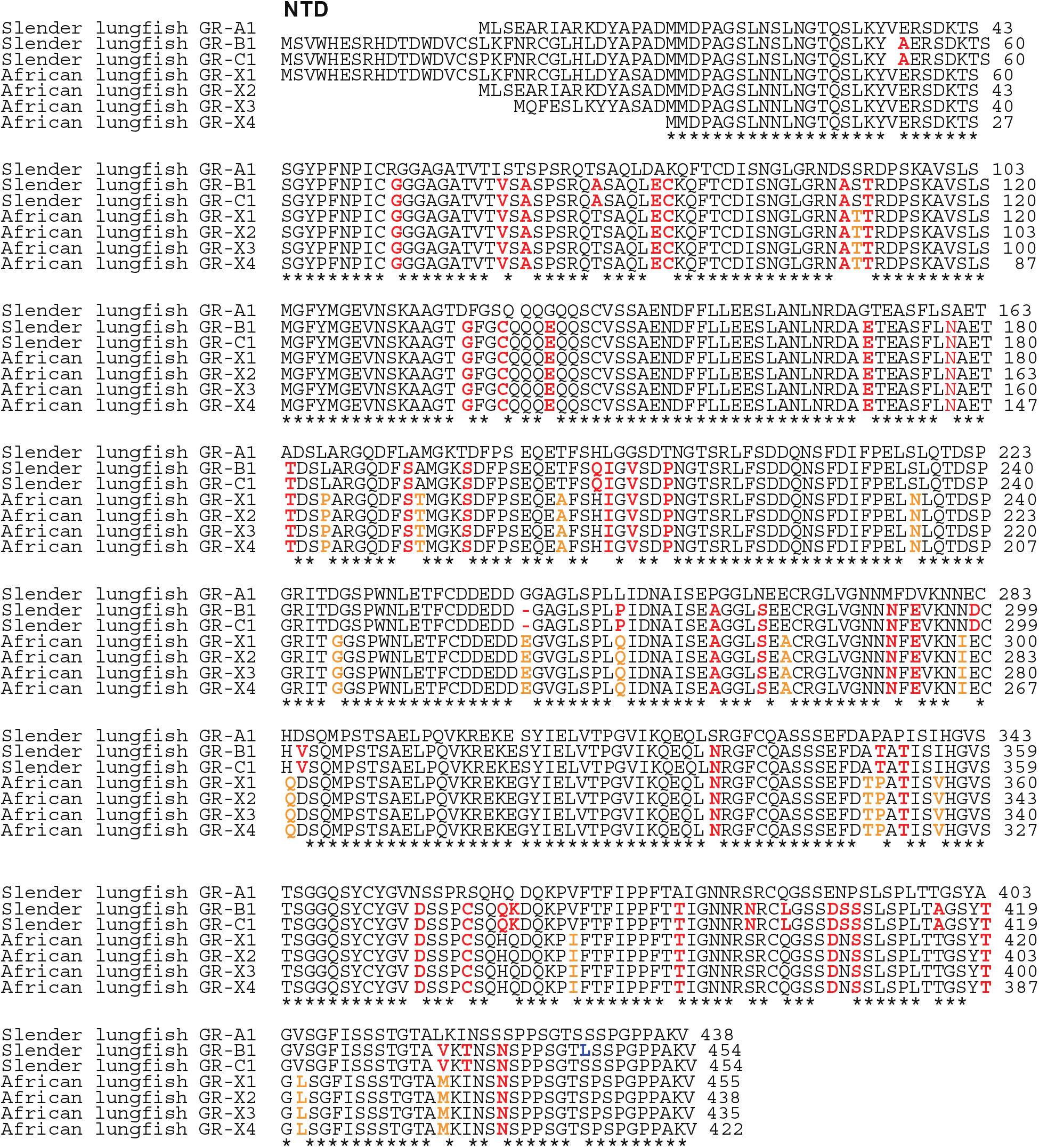
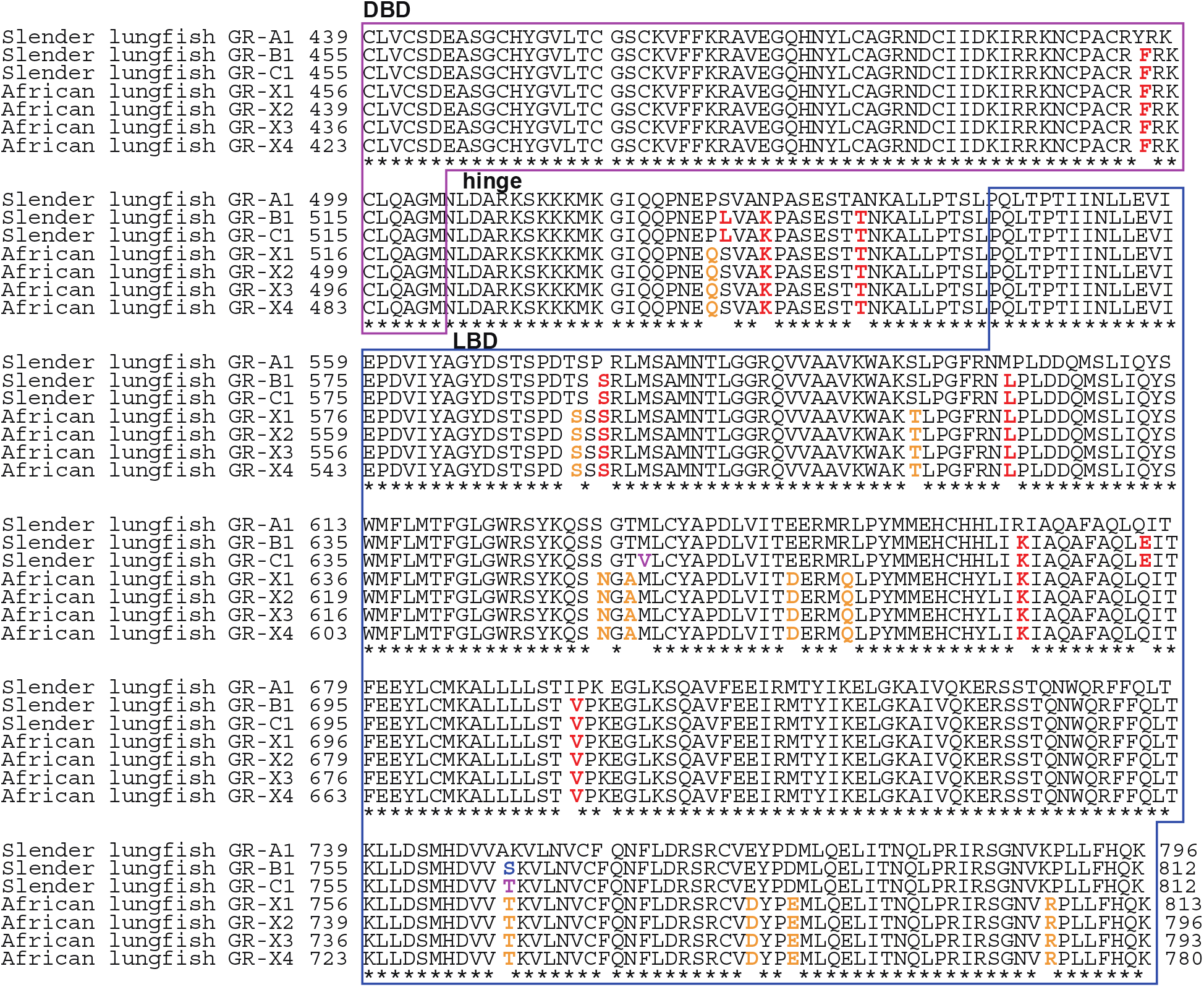
Multiple Alignment of the amino acid sequences of three African lungfish GRs and four West African lungfish GRs. West African lungfish glucocorticoid receptor sequences were downloaded from GenBank (Accessions XP_043925084 for X1, XP_043925085 for X2, XP_043925087 for X3, XP_043925088 for X4). Sequences were aligned with Clustal W [35], as described in the Methods section.

### 2C. Comparison of the amino acid sequences of slender African lungfish GR, West African lungfish GR, coelacanth GR, zebrafish GR and human GR

To begin to understand the relationship of lungfish GRs to other selected GRs, we constructed a multiple sequence alignment of slender African lungfish GR with West African lungfish GR, coelacanth GR, zebrafish GR and human GR (Figure 3). The DBD and hinge domains are highly conserved in all GRs. There is good sequence conservation of the LBD in all six GRs. However, there is an interesting pattern of sequence conservation in the NTD. There is excellent sequence conservation in the NTD among slender African lungfish GR, West African lungfish GR, coelacanth GR and human GR. The stronger conservation of the NTD in lungfish GRs with human GR than with zebrafish GR, indicates that the NTD in zebrafish GR has diverged from the other GRs.

**Figure 3.**
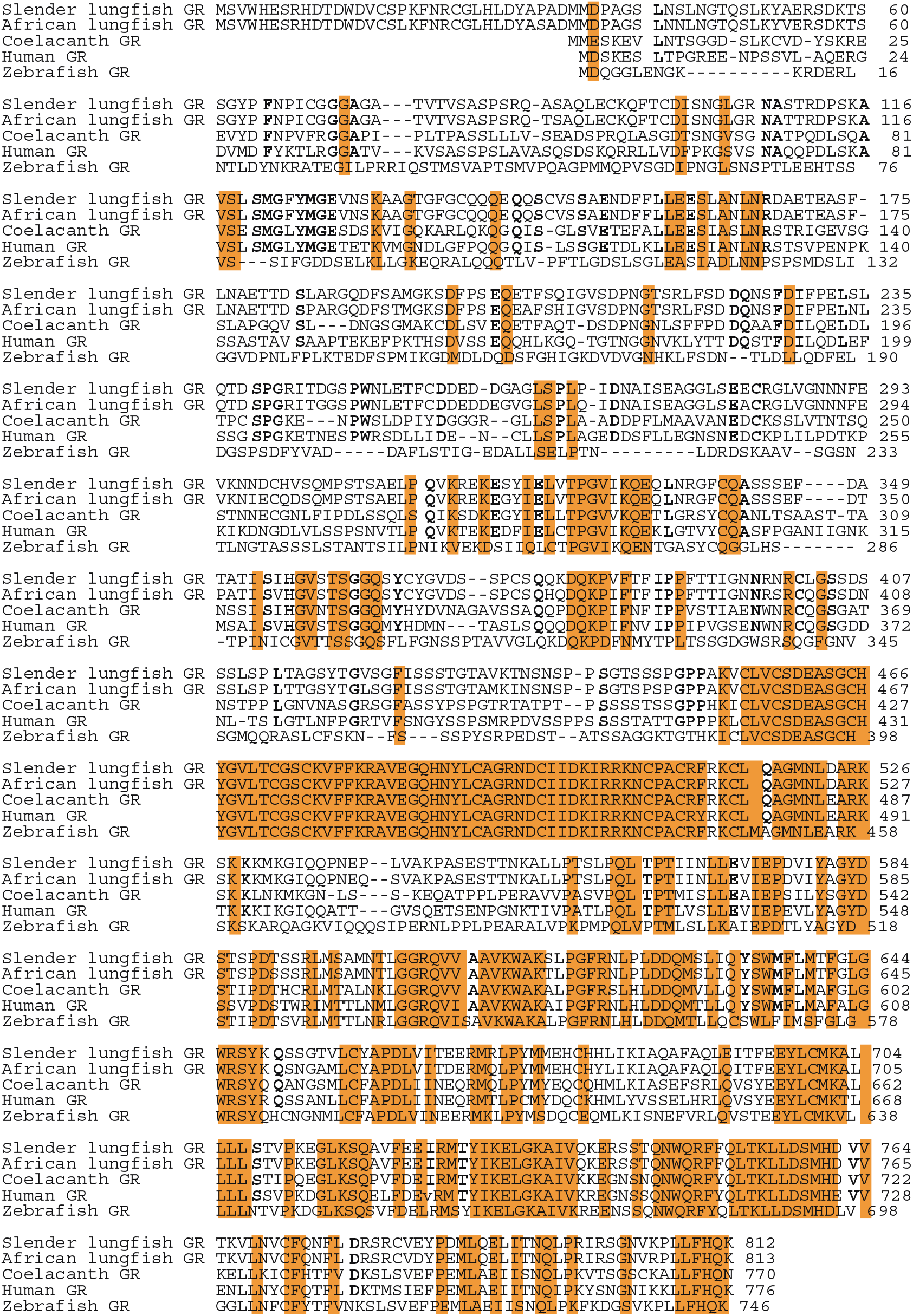
Multiple Sequence Alignment of slender African lungfish GR, West African lungfish GR, coelacanth GR, zebrafish GR and human GR. Glucocorticoid receptor sequences were downloaded from GenBank (Accession no. NP_000167 for human GR, XP_005996162 for coelacanth GR, and NP_001018547 for zebrafish GR) and aligned with Clustal W [35], as described in the Methods section. The NTD in zebrafish GR has gaps and sequence differences with the other GRs.

### 2D. Comparison of functional domains in slender African lungfish GR with domains in West African lungfish GR, coelacanth GR, zebrafish GR and human GR

Figure 4 shows the percent identity in the comparison of the different functional domains on slender African lungfish GR with the GR and MR from other vertebrates.

**Figure 4.**
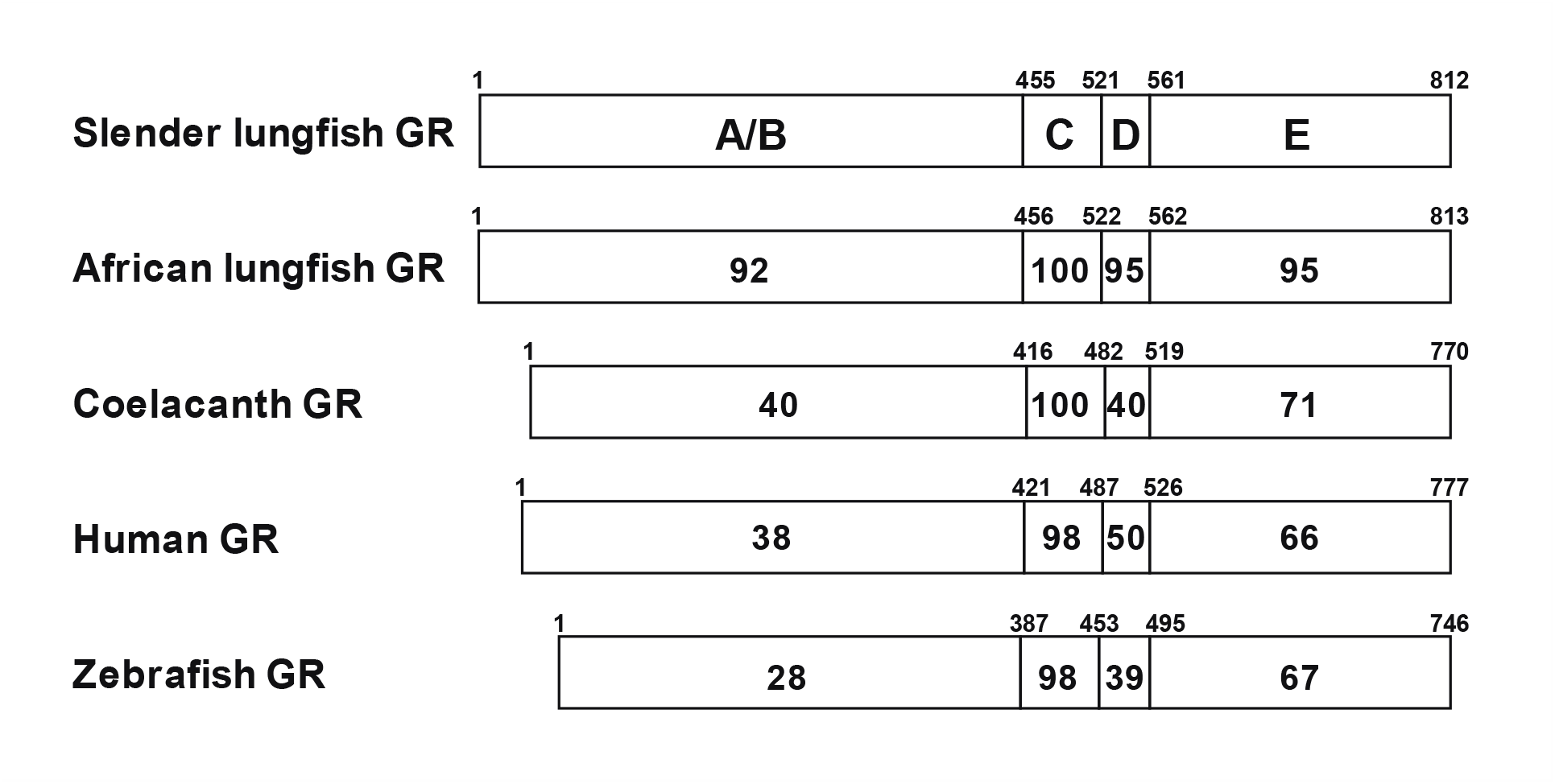
Comparison of functional domains of slender lungfish GR with domains in West African lungfish GR, coelacanth GR, zebrafish GR, human GR. Comparison of domains in slender African lungfish GR with GRs from West African lungfish, coelacanths, humans and zebrafish and MRs from slender African lungfish, West African lungfish, humans and zebrafish. The functional NTD (A/B), DBD (C), hinge (D) and LBD (E) domains are schematically represented with the numbers of amino acid residues and the percentage of amino acid identity depicted.

As shown in figure 4, the DBD and LBD are highly conserved in all GRs. For example, slender African lungfish GR and human GR have 98% and 66% identity in DBD and LBD, respectively. There are similar % identities between corresponding DBDs and LBDs in lungfish GR and other GRs. This strong conservation of the DBD and LBD contrasts with the lower sequence identity between the NTD of slender African lungfish GR and human GR (38%) and even lower sequence identity with the NTD in zebrafish GR (28%).

### 2E. Phylogenetic Analysis

To better understand the relationships among the nine *P. dolloi* GRs and four *P. annectens* GRs, we constructed the phylogenetic tree, shown in Figures 5. In this phylogeny, the four African lungfish GRs cluster into one group. Slender African lungfish GR-A1 and GR-A2 are in a separate branch from the other slender African lungfish GRs. GR-A2 appears to be formed by alternative splicing of GR-A1. GR-B1, GR-B2 and GR-B3 cluster. GR-C3 and GR-C4 cluster, and GR-C4 appears to be formed by alternative splicing of GR-C3.

**Figure 5.**
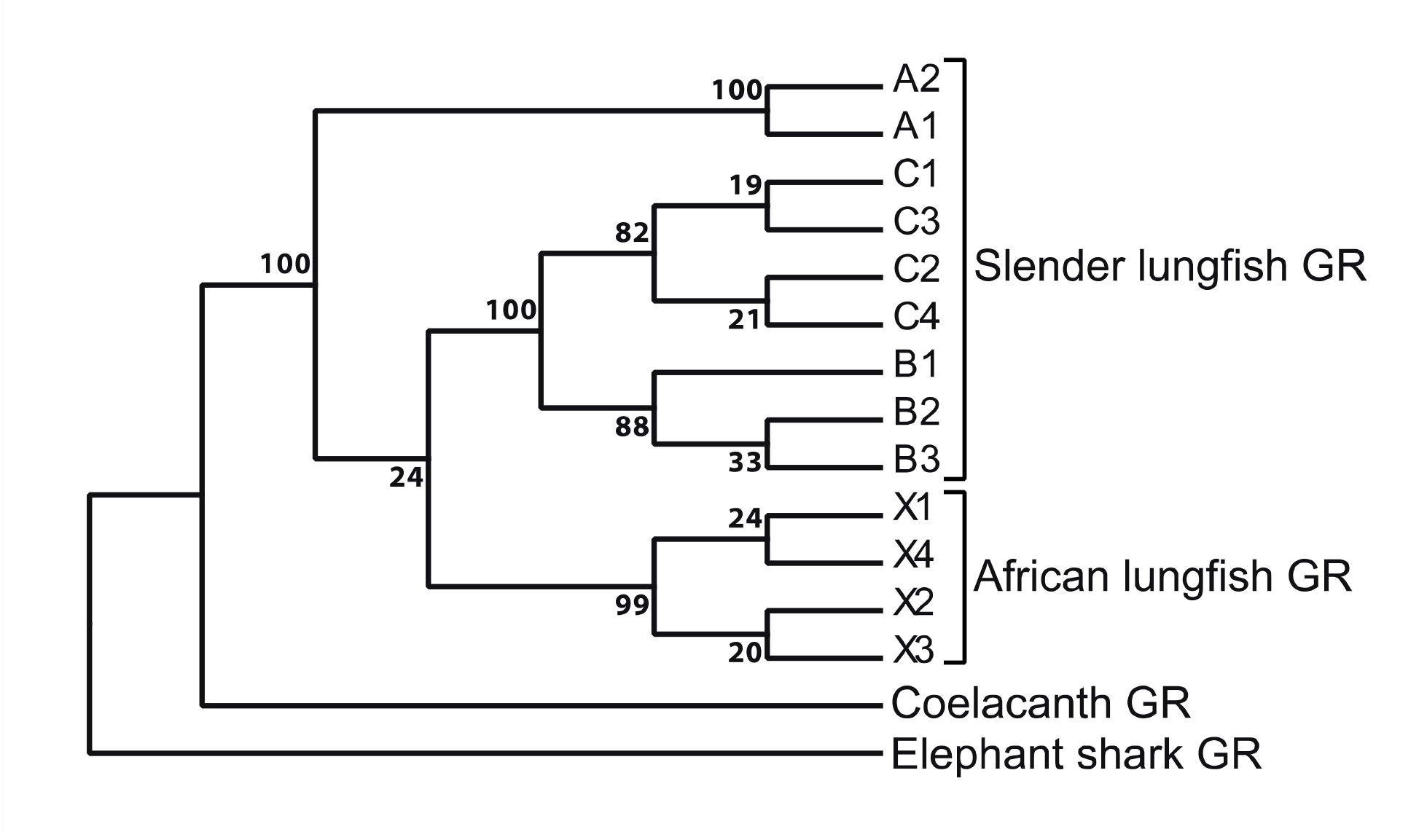
Phylogeny of slender African lungfish Glucocorticoid Receptors, West African lungfish Glucocorticoid Receptors, Coelacanth GR and Elephant Shark GR. MEGA5 [36] was used to construct this phylogeny. Statistics are based on 1,000 runs.

### 2F. Basis for the failure to clone *P. dolloi* GR

Figure 6 shows the location of the PCR primers that we used to successfully clone GRs from chicken, alligator and frog [31]. Due to the strong conservation of the GR and MR these PCR primers retrieved partial sequences from both the GR and MR in chicken, alligator and frog. The full sequences of these GRs and MRs was achieved in the next step using RACE. Our failure to clone *P. dolloi* GR was due using WQRF**Y**Q instead of WQRF**F**Q for the 1^st^/2^nd^-reverse primer. When we used WQRF**F**Q we were able to clone *P. dolloi* GR.

**Figure 6.**
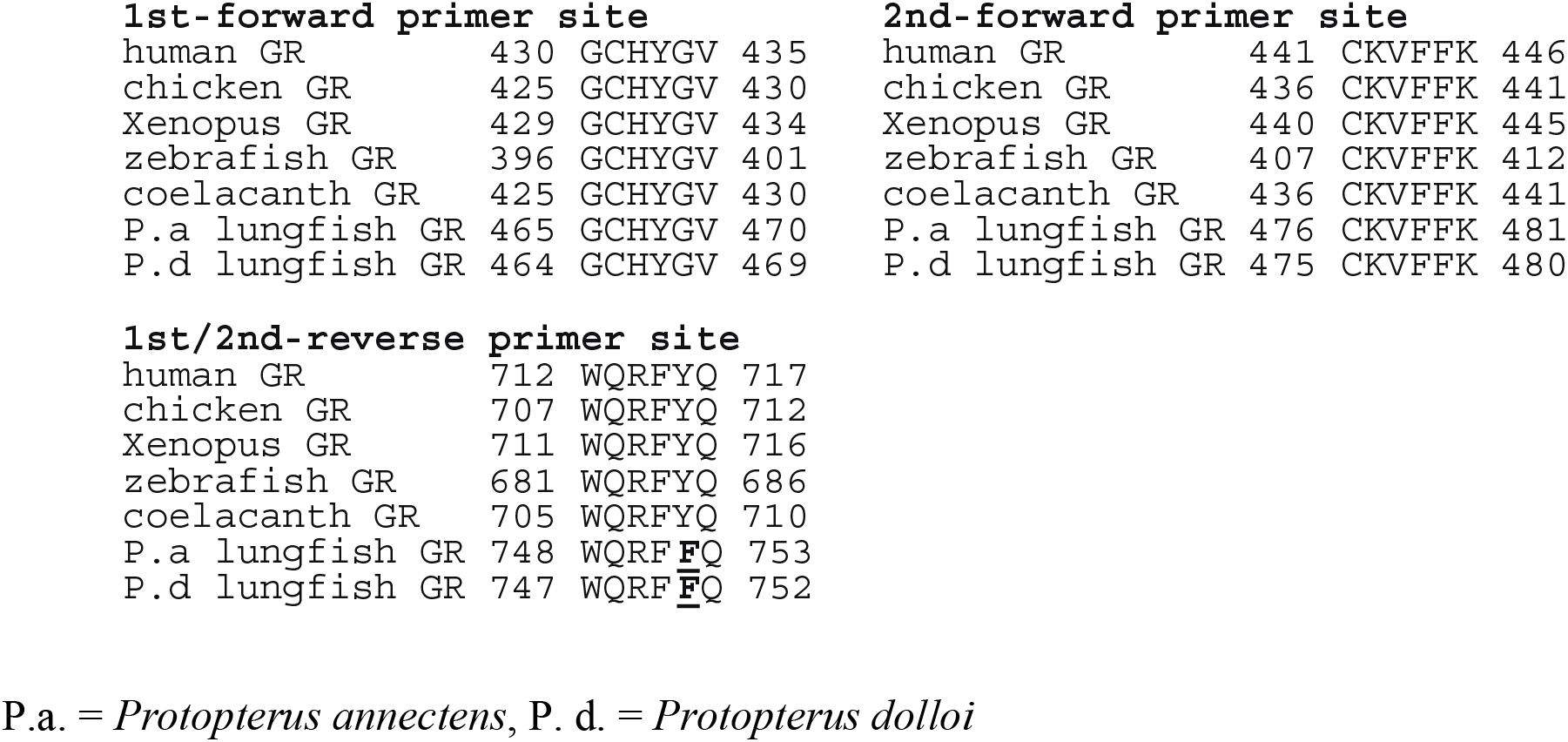
Location of PCR primers used for cloning of slender African lungfish GR, coelacanth GR, elephant shark GR, zebrafish GR and human GR. The correct 1^st^/2^nd^-reverse primer for PCR cloning of *P. dolloi* GR is WQRF**F**Q instead of WQRF**Y**Q.

### 2G. Summary

*P. dolloi* contains nine GR isoforms, in contrast to *P. annectens*, which contains four GR isoforms. We do not know how many GR isoforms are in Australian lungfish (*Neoceratodus forsteri*) because their GR sequences have not been deposited in GenBank. The availability of sequences of *P. dolloi* GRs and *P. annectens* GRs should permit using PCR to clone *N. forsteri* GRs, which would elucidate the number GR isoforms in this lungfish and the relationship of their GRs to the GRs of *P. dolloi* and *P. annectens*.

The response to corticosteroids of any lungfish GR is not known, nor are the functions of the multiple GR isoforms in *P. dolloi* GRs and *P. annectens* GRs. We have initiated studies to determine corticosteroid activation of *P. dolloi* GRs to begin to elucidate the functions of slender African lungfish GRs. It is interesting that there are multiple isoforms of human GR, due to alternative splicing of human GR, and these isoforms are important in achieving functional diversity of human GR [6,8,34,37]. A similar scenario is likely for *P. dolloi* GRs and *P. annectens* GRs.

## 3. Methods

### 3A. Animal

A slender spotted African lungfish, *Protopterus dolloi,* was purchased from a local commercial supplier. Lungfish were anesthetized in freshwater containing 0.02% ethyl 3-aminobenzoate methane-sulfonate from Sigma-Aldrich, and tissue samples were quickly dissected and frozen in liquid nitrogen. Animal handling procedures conformed to the guidelines set forth by the Institutional Animal Care and Use Committee at the University of Tokyo.

### 3B. Molecular cloning of lungfish *P. dolloi* glucocorticoid receptor

For P. dolloi GR cloning, we designed 4 types of forward N-terminal primers: F-X1: 5’-GTCATTTTCCCCGTGCTTAACGAA-3’, F-X2: 5’-GTCTGCAGCTTGAAACTTTGTAAC-3’, F-X3: 5’-GACGAACATGCTGACCGGATCATAA-3’, and F-X4: 5’-CATACTGCATTTACCAGAATAGAC-3’ and one C-terminal Reverse primer: R: 5’-GTTAAGGCAAATTTCTGATATTAAGGCAG-3’ based on the sequences of *P. annectens* GR (X1: XM_044069149, X2: XM_044069150, X3: XM_044069152, X4: XM_044069153). PCR was performed using four primer sets (F-X1xR, F-X2xR, F-X3xR, and F-X4xR) with ovary cDNA of *P. dolloi*, and the amplified DNA fragments with KOD-plus-DNA polymerase were subcloned into a cloning vector, pCR-BluntII-TOPO, and sequence analysis was performed for 10 or more clones for each primer sets.

### 3C. Database and sequence analysis

GRs for phylogenetic analysis were collected with Blast searches of Genbank. A phylogenetic tree for GRs was constructed by Maximum Likelihood analysis based on the JTT + G model after sequences were aligned by Clustal W [35]. Statistical confidence for each branch in the tree was evaluated by the bootstrap methods [38] with 1000 replications. Evolutionary analyses were conducted in MEGA5 program [36].

## 4. AUTHOR CONTRIBUTIONS

Y.K., S.O., L.B. and M.E.B. carried out the research and analyzed data. S.H. aided in the collection of animals. X.L. constructed plasmid DNAs used in this study. Y.K. and M.E.B. conceived and designed the experiments. Y.K., L.B. and M.E.B. wrote the paper. All authors gave final approval for publication.

